# Chromosomal Restructuring and Subgenome Divergence Drive Post-Polyploid Adaptive Diversification in *Sinocyclocheilus* Cavefish

**DOI:** 10.1101/2025.09.22.677718

**Authors:** Tingru Mao, Yewei Liu, Hannes Svardal, Mariana Vasconcellos, Jian Yang, Liandong Yang, Shunping He, Madhava Meegaskumbura

**Affiliations:** Guangxi Key Laboratory for Forest Ecology and Conservation, College of Forestry, Guangxi University, Nanning, Guangxi, PR China; The Key Laboratory of Aquatic Biodiversity and Conservation of Chinese Academy of Sciences, Institute of Hydrobiology, Chinese Academy of Sciences, Wuhan, 430072, PR China; Evolutionary Ecology Group, Department of Biology, University of Antwerp, Antwerp, Belgium; Understanding Evolution group, Naturalis Biodiversity Center, Leiden, The Netherlands; Departamento de Zoologia, Instituto de Biociências, Universidade de São Paulo, São Paulo, Brazil; Key Laboratory of Environment Change and Resource Use, Beibu Gulf, Nanning Normal University, Nanning, Guangxi, PR China

**Keywords:** Allopolyploidy, Subgenome Dominance, Adaptive Diversification, Cave Adaptation, Transposable Elements, Epigenetic Regulation

## Abstract

Allopolyploidy generates extensive genomic potential, yet its role in adaptive diversification remains poorly understood. We assembled phased chromosome-level genomes of cave and surface ecotypes of *Sinocyclocheilus*, a rapidly diversifying cavefish lineage, the most species-rich in the world. Our analyses show that all species derive from the ancestral allotetraploidisation event common to all cyprinids without subsequent polyploidisation. Instead, diversification has been driven by chromosomal rearrangements and the emergence of subgenome asymmetry. The dominant D subgenome maintains more genes, exhibits higher expression, and is enriched for adaptive functions, while the submissive S subgenome is inclined towards housekeeping roles. This asymmetry is further reinforced by the accumulation of transposable elements near genes exhibiting expression biased toward the S-subgenome. We also identify a derived epigenetic mechanism in the cave ecotype, i.e., differential methylation of these elements, absent in the surface ecotype. These findings show how structural, regulatory, and epigenetic processes have interacted following allopolyploidy to produce ecological and phenotypic diversity in *Sinocyclocheilus*. Our findings also offer quantitative evidence that cave adaptation is a complex process characterised by an evolutionary trade-off – the regression of energetically expensive traits combined with the remodeling of neural and developmental pathways driven by natural selection.

## INTRODUCTION

Whole-genome duplication (WGD) has long been recognised as a key driver of evolution, offering extra genetic material that can give rise to new functions and regulatory patterns (*1–4*). In polyploid organisms, duplicated gene sets, or subgenomes, often diverge over time in terms of structure and function. Genes may be lost, undergo subfunctionalisation, or develop new functions; such divergence can create long-term asymmetries in gene content, expression, and regulation (*5*, *6*). These asymmetries are increasingly linked to evolutionary innovation. However, their mechanistic connections to adaptive phenotypes remain poorly understood, especially under extreme environmental pressures.

Cave habitats and associated fauna offer an ideal backdrop to explore this problem. These ecosystems are characterised by constant darkness, low productivity, and relative environmental stability (*7*). Such conditions apply consistent selection pressures that often lead to convergent regressive traits, such as eye degeneration and loss of pigmentation, as well as constructive changes like the improvement of mechanosensory and chemosensory systems and the reorganisation of metabolic pathways (*8–11*). While the genetic foundations of these changes have been examined in diploid cave taxa (*12*), the behaviour of polyploid genomes under similar conditions remains mostly unexplored. Understanding how each subgenome influences phenotypes requires phased, chromosome-level assemblies. Such resources have hitherto been unavailable for cave ecotype polyploid vertebrates, hindering a full understanding of polyploid genome evolution in this context.

Adaptation to the extreme cave environment is a classic example of evolution in action, involving a complex interplay of regressive and constructive processes (*8*). Regressive evolution, initiated by the relaxation of purifying selection on functions like vision and pigmentation that are costly to maintain and useless in darkness, is further driven by positive selection for energy conservation (*9*, *10*, *13*, *14*). This adaptive trade-off is considered a highly efficient mechanism for shaping the cave phenotype (*12*, *15*). Concurrently, constructive evolution reshapes other systems for survival in perpetually dark environments. These changes include the enhancement of non-visual sensory modalities, such as mechanosensation and chemosensation, and the remodeling of metabolic pathways to maintain homeostasis despite scarce and intermittent food supplies (*10*, *11*, *14–16*). However, it remains unknown whether the genetic basis of these constructive traits has been subject to positive selection. The genetic architecture of these adaptive trade-offs offers critical insights into the core principles of evolutionary change.

An excellent example for studying these processes is the freshwater fish genus *Sinocyclocheilus*, which represents the largest cave-associated diversification of a vertebrate group (*17*). Native to the karst regions of Southwest China, this genus includes more than 80 species that exhibit a remarkable range of forms, from surface-dwelling species to fully subterranean species (*18*). Cave ecotype species show classic stygomorphic traits, such as eye degeneration, loss of pigmentation, and enhanced non-visual sensory systems. This extraordinary diversity, which developed over a relatively short evolutionary timescale, raises a vital question: what genomic factors have driven such a remarkable adaptive diversification? Whole-genome duplication, or polyploidization, serves as a powerful driver of evolutionary innovation by generating large pools of duplicate genes that can be shaped by natural selection (*1*, *2*). This process is recognised as a key factor in diversification within the tree of life, including in vertebrates (*4*, *19*). For instance, teleost fishes share an ancient duplication known as the teleost-specific third round of whole-genome duplication, TS3R (*20*, *21*). Subsequently, multiple lineages have undergone additional independent genome duplications, including species in the orders Cypriniformes (e.g., the common carp *Cyprinus carpio* and goldfish *Carassius auratus*) (*22*, *23*) and Salmoniformes (*24*, *25*). Similarly, *Sinocyclocheilus* is polyploid (*26*), yet its recent and rapid diversification remains somewhat mysterious. Did this diversification stem from the latent potential of the ancient TS3R, or was it driven by subsequent lineage-specific polyploidy events providing the necessary genomic catalyst?

Following allopolyploidisation, which involves genome duplication through interspecific hybridisation, there is often a phase of intense genomic restructuring. As the combined genomes settle into a stable new state, they undergo diploidisation, commonly characterised by large-scale chromosomal rearrangements (*27*). Processes such as species-specific chromosome fusions and fissions can create reproductive barriers, accelerating the process of speciation (*28*, *29*). The genus *Sinocyclocheilus* exhibits these distinctive genomic features, with different species possessing varying chromosome numbers (*30–32*). This characteristic makes *Sinocyclocheilus* an excellent model system for investigating how large-scale chromosomal rearrangements and subsequent diploidisation contribute to speciation following polyploidisation.

The presence of two distinct parental subgenomes within a single nucleus leads to a unique evolutionary process. Often, one subgenome becomes dominant, as noted by Bird et al. (*33*). This dominant subgenome generally preserves more genes, exhibits higher expression levels, and undergoes stronger purifying selection. Conversely, the recessive subgenome has more freedom to develop new functions, as discussed by Alger & Edger and Cheng et al. (*5*, *6*). This division of roles may offer benefits: one subgenome can focus on essential cellular functions, while the other evolves to meet new ecological challenges. However, such subgenome dynamics have remained largely unexplored in allotetraploid fish of the genus *Sinocyclocheilus*.

While the patterns of gene loss and selection are now better understood, the regulatory mechanisms driving adaptive changes in complex polyploid genomes are still not fully clear. Gene expression results from interactions between genetic and epigenetic components (*34*). Additionally, transposable elements are increasingly acknowledged as important sources of regulatory innovation (*35*, *36*). Epigenetic modifications, including DNA methylation, and the three-dimensional organisation of chromatin, contribute significantly to gene regulation. (*37*). The manner in which these landscapes evolve following whole-genome duplication, and their role in subgenome dominance and adaptation to extreme environments such as caves, remains poorly explored, particularly in vertebrates. The freshwater fish genus *Sinocyclocheilus* offers an ideal natural experiment to tackle these questions.

We generated high-quality, chromosome-level genome assemblies for two representative *Sinocyclocheilus* species: the cave-adapted *S. tianlinensis* (clade E) and the surface-dwelling *S. longibarbatus* (clade B). These assemblies were analysed together with six publicly available genomes from other major clades to enable a broad comparative genomic and epigenomic study. By integrating chromosome-scale assemblies with multi-tissue transcriptomes and whole-genome bisulfite sequencing, we investigate the evolutionary processes shaping this genus.

Our genomic analysis of the cave ecotype of *Sinocyclocheilus* reveals the complex interplay of selective forces that have shaped its adaptation to a subterranean existence. We track their shared polyploid origin, evaluate the impact of large-scale chromosomal rearrangements on diversification, and examine the structural, transcriptional, and functional asymmetries between the ancestral subgenomes. Additionally, we analyse regulatory mechanisms such as transposable elements, DNA methylation, and three-dimensional chromatin organisation that maintain subgenome dominance. These processes are linked to differences between cave and surface ecotypes.

## RESULTS AND DISCUSSION

### Karyotype and Genome Size Estimation

To determine the karyotypes of *S. tianlinensis* and *S. longibarbatus*, we assessed chromosome preparations from approximately 50 metaphase spreads for each species (Supplementary Fig. 1). Our examination of mitotic metaphase cells in *S. tianlinensis* revealed chromosome counts ranging from 92 to 104. A clear modal number of 100 was recorded in 60% of cells (30 out of 50). Other observed counts were 96 (10 cells), 98 (5 cells), 104 (2 cells), and 92 (3 cells). On this basis, the somatic chromosome number was determined as 2n = 100. In *S. longibarbatus*, chromosome counts ranged from 94 to 102. The modal number of 100 occurred in 64% of spreads (32 out of 50), while other values included 96 (13 cells), 98 (3 cells), 102 (1 cell), and 94 (1 cell). The consistency of the modal value at 100 suggests this as the somatic chromosome number of both species. Deviations from the modal value in both taxa are most likely explained by aneuploidy or technical artefacts arising during slide preparation, such as chromosome loss.

To characterise the genomes of *S. tianlinensis* and *S. longibarbatus*, we performed k-mer analysis using Illumina sequencing data (Supplementary Fig. 1; Supplementary Table 2). For *S. tianlinensis*, we estimated the genome size to be 2.02 Gbp, with a repeat content of 76.62% and a heterozygosity rate of 0.18%. In the case of *S. longibarbatus*, we estimated a genome size of 1.59 Gbp, of which 70.02% consists of repeat sequences, and determined a heterozygosity rate of 0.38%.

### High-Quality Genome Assembly and Annotation

To elucidate the genomic architecture, we generated high-quality, chromosome-level genome assemblies for *S. tianlinensis* and *S. longibarbatus*. Our initial assemblies, based on HiFi reads, yielded a 2,004 Mb genome for *S. tianlinensis* (N50 = 25.72 Mb) and a 1,870 Mb genome for *S. longibarbatus* (N50 = 29.29 Mb). Following the initial assembly, we employed Hi-C scaffolding to organise both assemblies into 50 chromosome-level scaffolds. For *S. tianlinensis*, we successfully anchored 1,907 Mb (95.18%) of the sequence, achieving a scaffold N50 of 38.41 Mb. In the *S. longibarbatus* assembly, we anchored 1,868 Mb (99.85%) of the sequence, which resulted in a scaffold N50 of 37.38 Mb. We subsequently confirmed the accuracy of the chromosome-scale scaffolding through Hi-C contact heatmaps, which revealed strong interaction signals primarily within each chromosome.

We performed a comprehensive genomic analysis to assess the quality of our genome assemblies and investigate the evolutionary history of *S. tianlinensis* and *S. longibarbatus*. We generated high-quality genome assemblies for both species, which exhibited high completeness with BUSCO scores exceeding 98%. Our analysis revealed extensive gene duplication in both genomes, with approximately 70-73% of BUSCO genes being duplicated, a finding consistent with their tetraploid status (Supplementary Table 5). Furthermore, we identified 50 chromosomes in both species, suggesting a derivation from an ancestral diploid karyotype of n = 25, which is typical for cyprinids (*38*). These genomic features confirm a shared whole-genome duplication (WGD) event and establish these species as a valuable model for studying the long-term consequences of polyploidy, such as gene fractionation and functional divergence.

To characterise the repetitive element landscape within the genomes, we conducted a detailed analysis of their composition and distribution. Our investigation revealed that repetitive sequences constitute a significant portion of both genomes, accounting for 57.56% in *S. tianlinensis* and 59.08% in *S. longibarbatus* (Supplementary Table 6). We identified DNA transposons as the most abundant repeat class in both genomes, comprising 30.46% of the S. tianlinensis genome and 31.16% of the S. longibarbatus genome. Within this class, we found the TcMar-Tc1 superfamily to be the most prevalent (Supplementary Table 7). Furthermore, we identified other transposable elements, including Long Terminal Repeats (LTRs), which constituted 7.50% and 9.19% of the respective genomes. Long Interspersed Nuclear Elements (LINEs) accounted for 6.80% and 6.53%, and Short Interspersed Nuclear Elements (SINEs) made up 0.22% and 0.31% in *S. tianlinensis* and *S. longibarbatus*, respectively (Supplementary Fig. 4; Supplementary Table 6).

We annotated 48,455 and 47,963 protein-coding genes for *S. tianlinensis* and *S. longibarbatus*, respectively (Supplementary Table 8). The completeness of these gene sets was validated using BUSCO, which identified 98.70% and 98.80% of expected genes in *S. tianlinensis* and *S. longibarbatus*, respectively, indicating high-quality predictions (Supplementary Table 9). For functional annotation, homology searches against public databases annotated a high percentage of genes. In the *S. tianlinensis* and *S. longibarbatus* genomes, genes with significant hits were: 83.55% and 86.23% in SwissProt; 98.55% and 98.69% in NR; and 98.49% and 98.65% in TrEMBL, respectively. Similarly, protein domain analysis annotated 96.38% and 94.53% of genes in InterPro, and 93.82% and 99.16% in Emapper for the two species, respectively. Ultimately, 99.07% of all predicted genes in *S. tianlinensis* and 99.16% in *S. longibarbatus* were successfully assigned a putative function in at least one of the queried databases (Supplementary Table 10).

### Phylogenomics and Gene Family Evolution

To resolve the evolutionary relationships among *Sinocyclocheilus* species and place our target species in a robust phylogenetic framework, we constructed a phylogenetic tree using a dataset of 3,710 single-copy orthologous genes from eight *Sinocyclocheilus* species (*S. jii*, *S. tianlinensis*, *S. rhinocerous*, *S. anshuiensis*, *S. longibarbatus*, *S. grahami*, *S. maitianheenis*, and *S. anophthalmus*) and two outgroup species (*Cyprinus carpio* and *Carassius auratus*). This analysis successfully recovered the six expected clades within the *Sinocyclocheilus* genus (Fig. 1b). In a broader comparative genomic analysis, we clustered all 420,131 protein-coding genes from these ten species into 47,295 orthologous groups (orthogroups). This included 7,475 core orthogroups present in all ten species and 1,347 species-specific orthogroups (Supplementary Table 11). Focusing on the target genus, we identified 12,541 gene families that were shared among all eight *Sinocyclocheilus* species (Fig.1d; Supplementary Table 12).

**Fig. 1.**
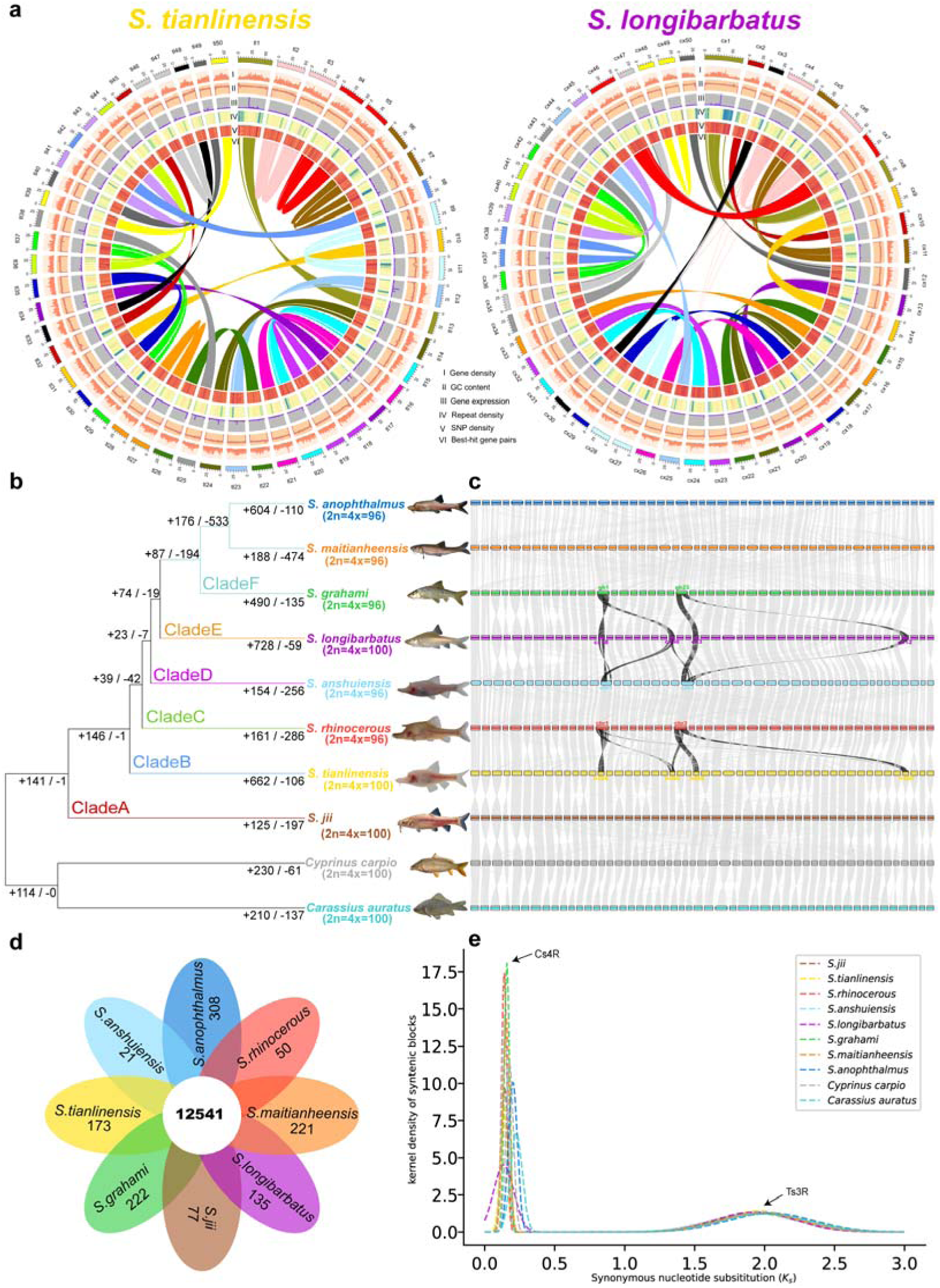
Evolution of the *Sinocyclocheilus* genome. (a) Features of homoeologous chromosomes in the *S. tianlinensis* and *S. longibarbatus* genomes. From outer to inner: I, gene density; II, GC content; III, gene expression based on muscle tissue; IV, repeat density; V, SNP density; VI, collinearity of syntenic genes. (b) A species tree resolving the phylogenetic relationships among eight *Sinocyclocheilus* species and two outgroups (*C. carpio* and *C. auratus*). The topology was inferred from 3,710 single-copy orthogroups using STAG and rooted via STRIDE. All nodes are maximally supported. Numbers on each branch quantify the count of significantly expanded (+) and contracted (−) gene families (P < 0.05). Chromosome numbers are provided in parentheses. (c) Chromosome-level macrosynteny among eight *Sinocyclocheilus* species, *C. carpio*, and *C. auratus*. Gray lines connect conserved syntenic blocks across the species. Black lines specifically highlight major chromosomal rearrangements (fusions and fissions) between key pairs: *S. tianlinensis* and *S. rhinocerous*; *S. anshuiensis* and *S. longibarbatus*; and *S. longibarbatus* and *S. grahami*. Chromosomes are grouped into subgenome S (first 24 or 25) and subgenome D (last 24 or 25). (d) Venn diagram of orthogroups in 8 *Sinocyclocheilus* genomes. The central number represents the core set of gene families all species share, while the numbers in the petals indicate the count of species-specific gene families. (e) Distribution of synonymous substitution rates (Ks) in 8 *Sinocyclocheilus* fishes, *C. carpio*, and *C. auratus*. The prominent peaks indicated by arrows, Ts3R (teleost-specific third-round whole-genome duplication) and Cs4R (cyprinid-specific fourth-round whole-genome duplication), provide evidence for ancient polyploidization events in these lineages.

To identify genomic signatures of convergent adaptation to cave environments, we analysed gene family evolution across the phylogeny. We focused on significant size changes on the terminal branches leading to each of the four cave ecotypes. This clade-specific analysis identified the number of inferred gene family expansions and contractions for *S. tianlinensis* (662 expanded, 106 contracted), *S. rhinocerous* (161, 286), *S. anshuiensis* (154, 256), and *S. anophthalmus* (604, 110), relative to their respective immediate ancestors (Fig. 1b).

To uncover evidence of parallel evolution, we performed a functional enrichment analysis on the intersection of gene families that were commonly expanded or contracted across these cave lineages (Supplementary Fig. 5; Supplementary Table 13). This approach isolates shared adaptive trends from lineage-specific changes. The set of commonly expanded gene families was significantly enriched in processes crucial for subterranean life, including enhancing nervous and sensory systems (e.g., chemosensory reception) and reconfiguring musculoskeletal systems. In stark contrast, the set of commonly contracted gene families was significantly enriched in functions related to the regression of vision-related pathways and simplification of the immune system. This strong signal of parallel evolution—where the same functional gene sets are repeatedly gained or lost in independent cave lineages—provides compelling evidence that these changes in gene family size seem to be driven by convergent selective pressures of the cave environment.

The genomic changes highlight the complex nature of cave adaptation, balancing the enhancement of essential sensory systems with the regression of functions that are redundant or energetically costly in dark environments. For instance, the significant enrichment of expanded gene families associated with nervous and sensory systems found in our analysis (Supplementary Fig. 5; Supplementary Table 13) points to a genetic basis for the heightened mechanosensory and chemosensory abilities often observed in cavefish (*11*). A notable example is the enhanced lateral line system in cavefish, which detects water movements and vibrations (*39*). Conversely, the significant contractions in gene families linked to vision-related pathways in our study (Supplementary Fig. 5; Supplementary Table 13) indicate an evolutionary trade-off, where the loss of vision-related genes conserves metabolic energy otherwise spent on maintaining a nonfunctional sensory system (*40*). Additionally, the contractions we observed in gene families tied to immune system specialisation (Supplementary Fig. 5; Supplementary Table 13) suggest adaptations to a distinct pathogen landscape in the relatively isolated cave environment, reducing the need for certain immune defences while specialising others (*41*). Similarly, alterations in reproductive processes through gene family contractions correspond to adaptations such as lower fecundity or modified breeding cycles, which are advantageous in ecosystems with scarce and unpredictable food resources (*42*).

### Shared Polyploidy and Chromosomal Reorganisation

To determine the timing and number of whole-genome duplication (WGD) events in the evolutionary history of *Sinocyclocheilus*, we analyzed the distributions of synonymous substitution rates (Ks) for paralogous gene pairs within all eight species, together with an outgroup including *Cyprinus carpio* and *Carassius auratus*. The Ks plots for all ten species consistently showed two distinct peaks: a recent peak with Ks values between 0.14 and 0.23, and an older peak between 1.92 and 2.04 (Fig. 1e). This bimodal distribution provides evidence that these species share two ancient whole-genome duplication (WGD) events. The recent peak corresponds to the carp-specific fourth-round WGD (CS4R) (*43*), while the older peak reflects the ancestral teleost-specific third-round WGD (TS3R) (*44*). Importantly, no additional peaks were detected in the *Sinocyclocheilus* species, indicating that they have not experienced lineage-specific WGDs. These results suggest that the extensive diversification of the genus was not driven by additional polyploidization events. Instead, we hypothesize that it was primarily facilitated by lineage-specific evolutionary trajectories of the duplicated genes derived from the CS4R, such as accelerated evolution, differential gene loss, and functional divergence.

Despite belonging to a single genus, *Sinocyclocheilus* species exhibit notable variation in chromosome number, pointing to a history of large-scale genomic reorganisation. Three species (*S. jii, S. tianlinensis, and S. longibarbatus*) retain a tetraploid karyotype of 2n = 4x = 100 (Supplementary Fig. 1) (*32*). In contrast, five species (*S. rhinocerous, S. anshuiensis, S. grahami*, *S. anophthalmus*, and *S. maitianheensis*) display a reduced chromosome number of 2n = 4x = 96 (*31*). Genome-wide syntenic comparisons confirmed the presence of extensive chromosomal rearrangements across the genus (Fig. 1c). We identified several key events, including chromosome fusions (e.g., in the common ancestor of *S. tianlinensis* and *S. rhinocerous*, and in the lineage leading to *S. grahami* after its divergence from *S. longibarbatus*) and chromosome fissions (e.g., in the ancestral lineage of the clade containing *S. anshuiensis* and *S. longibarbatus*). These events explain the observed variation in karyotypes.

Such rearrangements are characteristic of diploidisation (*27*) and can accelerate diversification by altering recombination patterns and promoting reproductive isolation (*45*). Analogous processes have been documented in the threespine stickleback (*46*), mudskipper (*47*), and swamp eel (*48*), where chromosomal restructuring has facilitated ecological adaptation. In *Sinocyclocheilus*, the CS4R WGD provided a broad genetic backdrop for adaptation, while subsequent karyotypic reorganisations likely accelerated cladogenesis, particularly under conditions of ecological isolation in cave systems. Therefore, these chromosomal shifts appear to be major evolutionary events that have facilitated speciation and adaptive diversification within the genus, rather than representing neutral variation.

### Positive and Relaxed Selection in Cave Ecotype Lineages

We examined signatures of selection in the cave ecotype *Sinocyclocheilus* by analysing 3,710 single-copy orthogroups with CODEML (positive selection) and RELAX (relaxed selection) (Supplementary Tables 14–15). This analysis identified 96 positively selected genes across 24 orthogroups, compared to 668 genes under relaxed selection within 167 orthogroups (Supplementary Table 16). The dominance of relaxed selection over positive selection provides quantitative support for the long-standing hypothesis that neutral, degenerative processes are a major driver of cave ecotype phenotypes (*49*).

Gene Ontology (GO) enrichment analysis of positively selected genes showed significant overrepresentation of terms associated with nervous system remodeling, morphogenesis, and signal transduction (Supplementary Fig. 6a; Supplementary Table 17). These results suggest that adaptation to aphotic environments involves regression and constructive evolutionary processes. A similar pattern has been documented in *Astyanax mexicanus*, where positively selected genes were enriched in developmental processes including embryogenesis, organogenesis, and cell morphogenesis (*50*). Conversely, genes under relaxed selection in *Sinocyclocheilus* were enriched for functions related to the degradation of vision and pigmentation and the restructuring of metabolism, circadian rhythms, and gene regulatory networks (Supplementary Fig.

6b; Supplementary Table 17). In the absence of light, the selective pressure required to maintain these energetically costly systems diminishes, leading to their gradual molecular decay (*49*, *51*). These highlight a dual process of regressive and constructive evolution that shapes cave adaptation. While relaxed selection leads to the regression of traits that are unnecessary in subterranean environments, positive selection also reshapes neural and developmental systems, creating a sensory toolkit vital for navigation, foraging, and survival in perpetual darkness.

### Allotetraploid Origin and Subgenome Identification

We applied the SubPhaser method to assign each chromosome of *S. tianlinensis* and *S. longibarbatus* to a subgenome (Figs. 2a, 2d). This approach relies on identifying subgenome-specific 15-mers, repetitive sequences that are highly enriched on one homoeologous chromosome relative to the other (*52*). The distinct distribution patterns of these markers enabled us to partition all chromosomes into two subgenomes, designated as ‘subS’ and ‘subD’ (Figs. 2b, 2c, 2e, 2f). The clear and consistent delineation of chromosomes into these two groups provides strong evidence for an allotetraploid origin in both species.

**Fig. 2.**
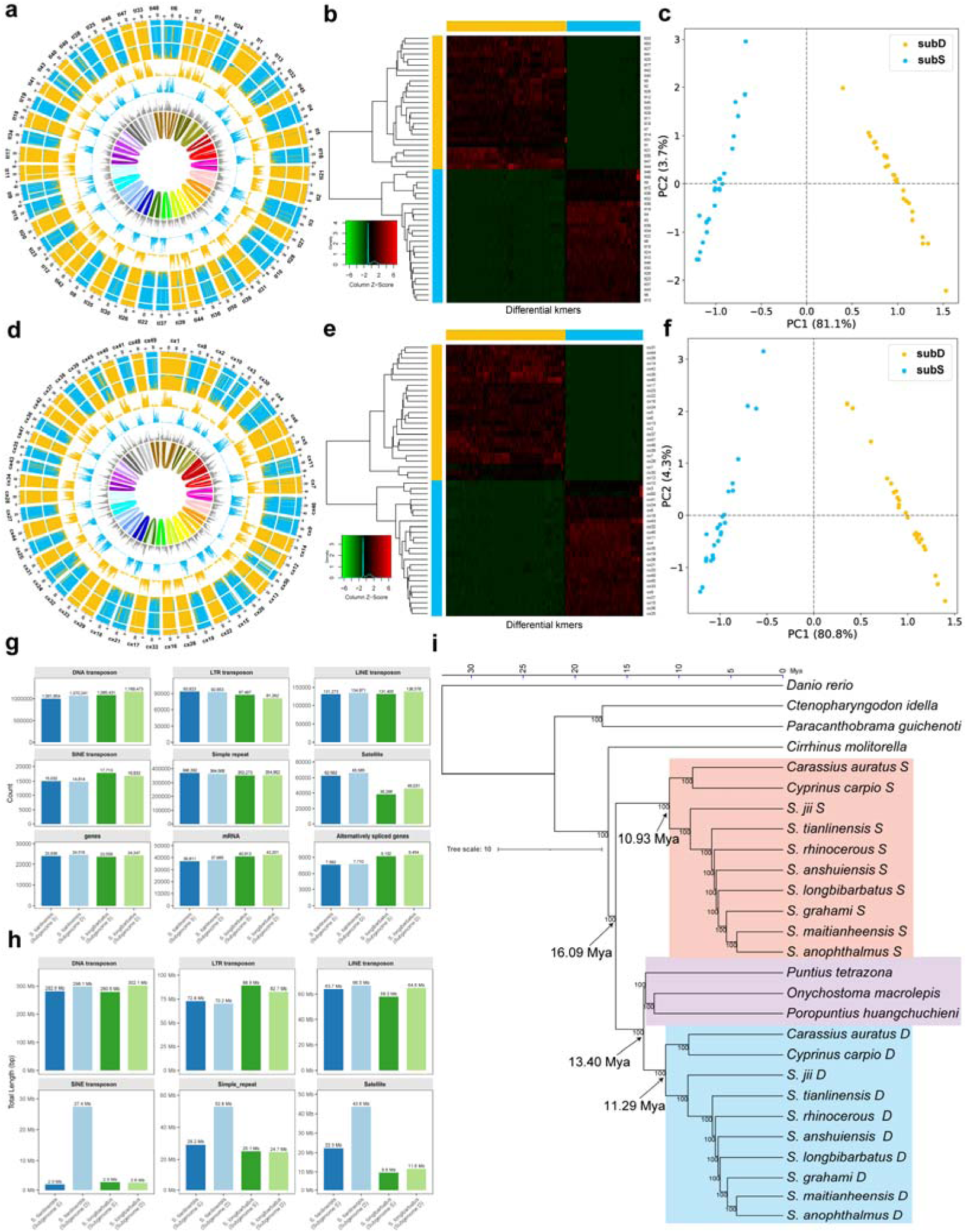
Allotetraploid origin and asymmetric evolution of subgenomes in *Sinocyclocheilus* fishes. (a) Chromosomal features of *S. tianlinensis* genome. Outer to inner circles: (1) subgenome assignments (k-means); (2) significant enrichment of subgenome-specific k-mers (color-coded; white: nonsignificant); (3) normalized proportion of subgenome-specific k-mers; (4–6) counts of subgenome-specific k-mers; (7) LTR-retrotransposon density (color indicates subgenome-specific enrichment; gray: nonspecific); (8) homoeologous blocks. Statistics computed in 1-Mb sliding windows. Homoeologous exchanges inferred. (b) Hierarchical clustering of *S. tianlinensis* chromosomes by differential k-mer frequencies, separating into two clades (subgenome D: yellow; S: blue). (c) PCA of k-mers validates distinct subgenome phasing. (d–f) Corresponding analyses for *S. longibarbatus* genome, showing similar chromosomal characteristics, clustering, and PCA separation. (g–h) Bar charts compare subgenome asymmetry, quantifying genomic feature counts (top) and lengths (bottom). (i) Maximum Likelihood phylogeny of cyprinid subgenomes and diploid relatives, inferred from 157 single-copy orthologs using IQ-TREE, with divergence times estimated by treePL. Includes subgenomes of eight *Sinocyclocheilus* species, *Cyprinus carpio*, *Carassius auratus*, and seven diploid cyprinids. Colored boxes denote clades: subgenome S (red), D (blue), and diploid sister lineage to D’s ancestor (purple).

To independently verify the allotetraploid origin of these species, we employed a complementary strategy based on comparative genomics and synonymous substitution rates (Ks). This approach is based on the principle that, in an allotetraploid, one subgenome will show greater similarity to a specific diploid outgroup, as indicated by lower Ks values. For this comparison, we selected the diploid tiger barb *(Puntius tetrazona*), a member of the Barbinae subfamily that is presumed to be closely related to one of the progenitor lineages (*53*). Using WGDI, we conducted a genome-wide syntenic analysis and calculated Ks values for homologous gene pairs between each tetraploid (*S. tianlinensis* and *S. longibarbatus*) and *P. tetrazona*.

As we expected, each tetraploid’s chromosomes separated into two sets when Ks values were plotted against the diploid genome. Chromosomes with consistently lower Ks values were assigned to one subgenome, while those with higher values were assigned to the other. Importantly, these Ks-based assignments were entirely consistent with the k-mer–based subgenome classification, providing robust, independent confirmation of the allotetraploid origin of both species (Supplementary Fig. 7).

### Evolution and Asymmetry of the Subgenomes

To further explore the effects of allotetraploidy, we compared the annotated content of the S and D subgenomes in both *S. tianlinensis* and *S. longibarbatus* (Figs. 2g, 2h; Supplementary Table 18). Our analysis showed a consistent pattern of asymmetric gene retention across both species. In each case, the D subgenome contained more annotated genes, mRNA transcripts, and alternatively spliced genes than the S subgenome. For example, in *S. tianlinensis*, the D subgenome had 24,516 genes compared with 23,939 in the S subgenome, a pattern also seen in *S. longibarbatus* (24,247 vs. 23,558) (Fig. 2g).

Patterns of repetitive element accumulation showed highly dynamic and lineage-specific processes. While DNA transposons were the most abundant class across all four subgenomes (Fig. 2g), striking differences were observed in the total sequence length of certain repeat families. In *S. tianlinensis*, the D subgenome showed dramatic expansions of SINEs (27.4 Mb), simple repeats (52.8 Mb), and satellites (43.6 Mb), all found in much smaller quantities in its S subgenome and in both subgenomes of *S. longibarbatus* (Fig. 2h). Conversely, the S subgenome of *S. longibarbatus* contained the largest proportion of LTR transposons (88.9 Mb). These findings suggest that asymmetric gene retention is a conserved feature of both species, whereas the amplification of repetitive elements has occurred in a lineage- and subgenome-specific manner following divergence.

To determine the evolutionary origin and timing of the WGD in *Sinocyclocheilus*, we built a Maximum Likelihood (ML) phylogeny using a concatenated dataset of 157 orthologous genes that are present as a single copy in each subgenome (Fig. 2i). The analysis included the subgenomes of ten tetraploid species (*Carassius auratus*, *Cyprinus carpio*, and eight *Sinocyclocheilus* species), along with several diploid cyprinid relatives. All major nodes achieved 100% bootstrap support, resulting in a highly robust topology. The resulting phylogeny offers clear evidence for a single, shared allopolyploid origin across these tetraploid lineages, consistent with previous findings for carp and goldfish (*23*, *43*).

The subgenomes of all tetraploid species formed two strongly supported monophyletic clades (Fig. 2i): an S subgenome clade (red) and a D subgenome clade (blue). Crucially, these were not recovered as sister groups. Instead, the D subgenome clade was sister to a lineage of diploid cyprinids (purple), including *P. tetrazona*, *Onychostoma macrolepis*, and *Poropuntius huangchuchieni*, with this entire group then sister to the S subgenome clade. This topology demonstrates that the WGD event arose through hybridisation between two diploid ancestors. Molecular dating places the hybridization event between 11 and 13 Mya, indicating that the WGD occurred prior to the major adaptive diversification of *Sinocyclocheilus* and likely provided the genetic framework for this radiation. This estimate is congruent with proposed dates for allopolyploidisation in related *Carassius* and *Cyprinus* lineages (*43*, *53–56*).

### Structural Variation and Homoeologous Exchange

To quantify genomic divergence between the subgenomes of *S. tianlinensis* and *S. longibarbatus*, we identified structural variations (SVs) using SyRI (Supplementary Fig. 8; Supplementary Table 19). This showed extensive chromosomal rearrangements in both species, including 279,592 SVs in *S. tianlinensis* and 273,802 in *S. longibarbatus*, composed mainly of translocations and duplications. Allelic comparisons uncovered millions of minor variants, comprising 4.88 million SNPs in *S. tianlinensis* and 4.80 million in *S. longibarbatus*.

Two consistent genomic patterns emerged. First, in both species, the D subgenome harboured a significantly greater number of gene duplications than the S subgenome (Supplementary Table 19). Gene duplication is widely recognised as a key driver of evolutionary novelty, providing substrates for neofunctionalisation and subfunctionalisation (*57*). This enrichment suggests that the D subgenome has acted as a primary locus of genetic innovation, aiding the emergence of new traits while maintaining ancestral functions. Second, the cave ecotype, *S. tianlinensis*, carried a greater number of SVs and SNPs than its surface ecotype counterpart, *S. longibarbatus*. This increased genomic divergence between the two species likely reflects a combination of demographic processes and the intense selective pressures typical of subterranean habitats. The scarcity of resources and absence of light in cave ecosystems impose profound morphological and physiological challenges (*58*). The elevated number of private mutations in *S. tianlinensis* likely stems from a combination of evolutionary processes characteristic of cave colonisation, though their relative contributions are difficult to disentangle. On one hand, non-adaptive processes likely played a significant role. Population bottlenecks during cave colonisation may have fixed numerous mutations through stochastic drift (*59*). Concurrently, the transition to darkness would have led to relaxed purifying selection on energetically costly traits that became redundant, such as vision and pigmentation, permitting the passive accumulation of mutations in these pathways. On the other hand, this expanded pool of genetic variation—comprising both neutral and potentially adaptive alleles—served as the raw material for positive selection. Extensive SVs, especially gene duplications, could have provided novel genetic substrates for constructive traits like enhanced sensory systems (*60*). Therefore, the genome of *S. tianlinensis* appears to be a mosaic, shaped jointly by the creative forces of directional selection acting on advantageous variation and the stochastic effects of genetic drift and relaxed selection prevalent in small, isolated cave populations—a dynamic also observed in other cave lineages such as Astyanax mexicanus (*61*).

Active interactions between subgenomes through homoeologous exchanges (HEs) have previously been documented in common carp (*43*, *53*). To investigate this process in *Sinocyclocheilus*, we analysed read-mapping depth to detect HEs between subgenomes. We identified 58 “HE with replacement” events in *S. tianlinensi*s, concentrated on the tl46–tl47 chromosome pair, and 248 events in *S. longibarbatus*, concentrated on the cx3–cx30 pair. The higher number of HEs in *S. longibarbatus* is likely due to its greater Illumina sequencing depth (∼54× versus ∼42× in *S. tianlinensis*), which increases detection sensitivity (Supplementary Table 2). While absolute counts are influenced by coverage, HEs in both species suggest ongoing genetic interaction between subgenomes. Despite millions of years of divergence, homologous chromosomes have not yet achieved complete meiotic isolation and seem to retain their capacity for exchange. Such persistent homology may allow the transfer of adaptive alleles and generate novel genetic combinations, contributing to long-term evolutionary flexibility.

### Mechanisms of Subgenome Dominance

To explore patterns of gene retention and loss following polyploidisation, we examined the distribution of single-copy BUSCO genes between the two subgenomes. Our analysis showed a consistent pattern of asymmetric gene loss, with subgenome D retaining significantly more BUSCO singletons than subgenome S in both *S. tianlinensis* and *S. longibarbatus* (Supplementary Fig. 9). In *S. tianlinensis*, subgenome D retained 3,101 complete single-copy BUSCO genes compared to 2,901 in subgenome S (χ² test; P < 0.001). Similarly, in *S. longibarbatus*, subgenome D retained 3,140 genes versus 2,928 in subgenome S (χ² test; P < 0.001). These results provide clear evidence of biased fractionation, with the S subgenome experiencing more extensive gene loss than the D subgenome after allopolyploidisation. This pattern establishes a definitive subgenome dominance hierarchy, consistent with the idea that the merging of two distinct genomes often results in preferential retention within one subgenome (*6*, *55*).

Subgenome dominance is also characterised by higher expression of genes from one subgenome relative to their homoeologous counterparts in the other (*6*). To test this, we used JCVI to identify homoeologous gene pairs between subgenomes D and S in *S. tianlinensis* and *S. longibarbatus*, and compared their expression across multiple tissues. Expression profiles were analysed in nine tissues (eye, skin, brain, liver, heart, gill, kidney, spleen, and muscle) for *S. tianlinensis* and eight tissues (eye, brain, liver, heart, gill, spleen, muscle, and swim bladder) for *S. longibarbatus* (Supplementary Tables 21–22). For all tissues, we calculated the log2-transformed expression ratio of homoeologous pairs (SubD TPM / SubS TPM). The resulting distributions were consistently skewed above zero (Figs. 3a–3i; 3n–u), indicating higher median gene expression from the D subgenome relative to the S subgenome across both species.

**Fig. 3.**
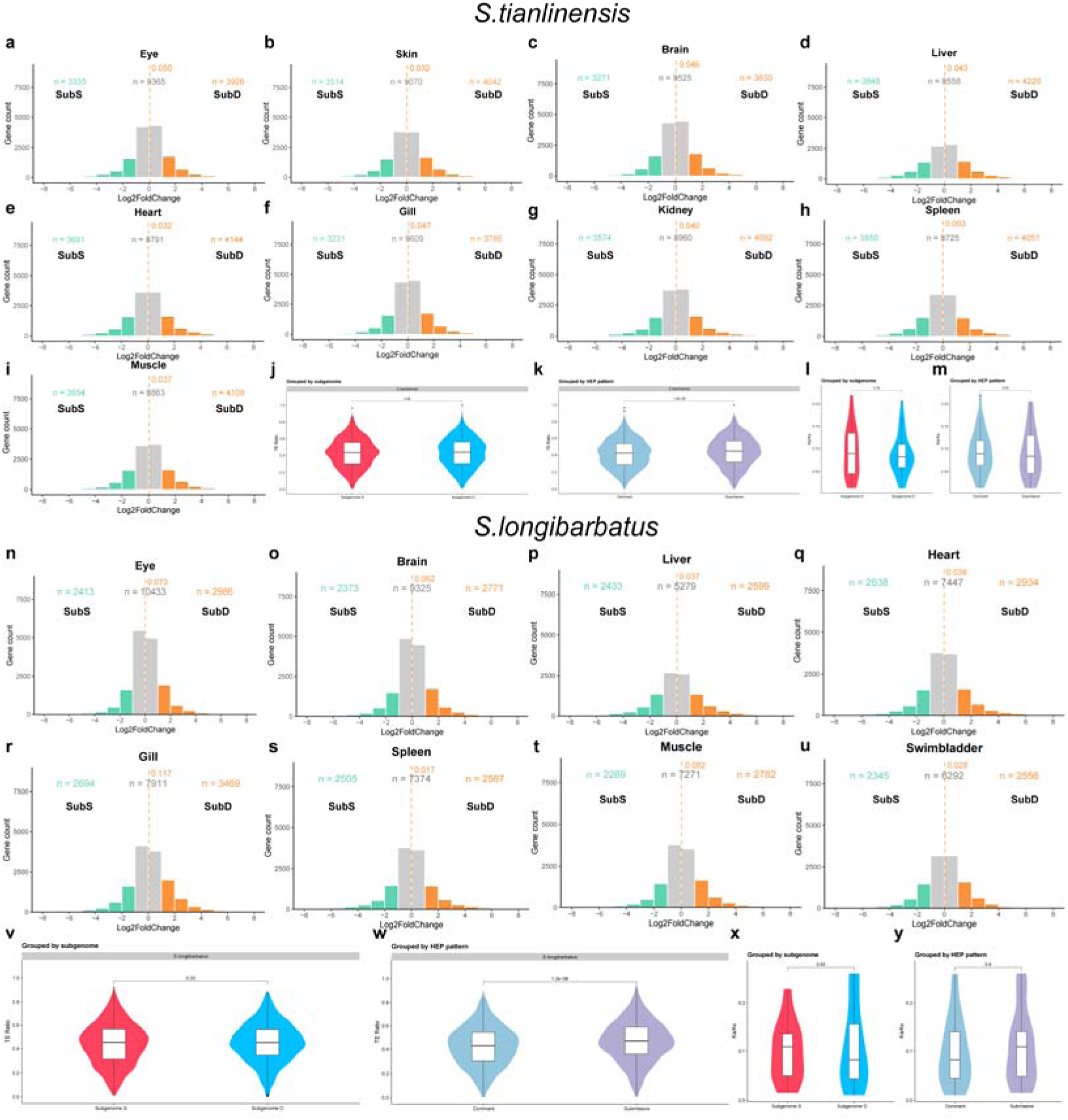
Biased homoeologous gene expression in *S. tianlinensis* and *S. longibarbatus*. (a–i) Histograms of homoeolog expression bias (HEB) in nine *S. tianlinensis* tissues (Eye, Skin, Brain, Liver, Heart, Gill, Kidney, Spleen, Muscle), showing log2 fold change (TPM subgenome D / TPM subgenome S). Genes are colored by expression pattern: subgenome S-biased (blue), D-biased (orange), or balanced (gray). HEB is prevalent across tissues. (j) Violin plot of TE ratios in flanking regions of *S. tianlinensis* genes shows no significant subgenome difference. (k) Dominantly expressed homoeologs in *S. tianlinensis* have lower flanking TE ratios than submissive ones (paired Wilcoxon test, P < 0.01), linking TE content to expression. (l) Violin plot of Ka/Ks ratios for *S. tianlinensis* homoeologs shows no significant subgenome difference. (m) No significant Ka/Ks difference between dominant and submissive homoeologs, indicating expression dominance is not tied to selective pressure. (n–u) HEB analysis in eight *S. longibarbatus* tissues (Eye, Brain, Liver, Heart, Gill, Spleen, Muscle, Swimbladder) shows similar patterns, with prevalent biased expression (S: blue; D: orange). (v) No significant TE ratio difference between subgenomes in *S. longibarbatus*. (w) Dominantly expressed homoeologs have lower TE ratios than submissive ones (P < 0.01). (x) No significant Ka/Ks difference between subgenomes. (y) No significant Ka/Ks difference between dominant and submissive homoeologs, consistent with *S. tianlinensis*.

These results demonstrate robust expression dominance of the D subgenome, a recurrent outcome of allopolyploidisation that has been well documented in plants such as cotton and Brassica (*62*, *63*), and also reported in vertebrates including the allotetraploid frog *Xenopus laevis* (*64*), common carp (*43*), and goldfish (*55*). By establishing *Sinocyclocheilus* within this paradigm, our findings provide a valuable vertebrate model for investigating the establishment, maintenance, and evolutionary consequences of subgenome dominance.

The proximity of transposable elements (TEs) to genes is known to influence their expression and has been linked to subgenome expression dominance in allopolyploid plants and fish (*54*, *65*). To examine this in *Sinocyclocheilus*, we evaluated TE density near homoeologous genes in both *S. tianlinensis* and *S. longibarbatus*. A comprehensive comparison of all genes in subgenomes D and S showed no significant overall bias in TE density in either species (Figs. 3j, 3v). However, a clear and significant pattern appeared when focusing on the subset of genes with consistent expression bias across all analysed tissues. In both species, dominantly expressed homoeologs (biased towards subgenome D) had notably lower densities of proximal TEs than their submissive counterparts (biased towards subgenome S) (Figs. 3k, 3w). This supports the idea that localised, gene-proximal TE landscapes, rather than global genome-wide trends, help establish and sustain subgenome dominance. These results align with previous research in bread wheat, brassicas, and other fish species (*54*, *66*, *67*), which have repeatedly connected lower TE presence near genes to subgenome dominance. Evidence from newly formed allopolyploids further shows that dominance can develop quickly, often alongside dynamic changes in TE methylation between dominant and recessive subgenomes (*68*). Our findings expand these observations by demonstrating that TE effects in *Sinocyclocheilus* are highly localised, indicating that the influence of TEs on gene regulation is concentrated in specific regulatory contexts rather than spread across the genome.

A second hypothesis suggests that expression asymmetry may arise from different selection pressures, with one subgenome experiencing relaxed constraints that allow functional divergence (*69*). To investigate this, we calculated Ka/Ks ratios for homoeologous gene pairs in *S. tianlinensis* and *S. longibarbatus*. No significant differences in selection pressure were found, either in overall comparisons of the D and S subgenomes (Figs. 3l, 3x) or between genes that are dominantly and submissively expressed (Figs. 3m, 3y). These findings align with observations in other cyprinid allopolyploids, such as goldfish and common carp, where subgenome dominance has been mainly linked to gene regulation rather than protein evolution (*53*, *55*). The lack of differential Ka/Ks ratios indicates that most homoeologous proteins are maintained under strong purifying selection, regardless of their expression levels.

This separation of regulatory and coding evolution allows for controlling gene dosage and expression, offering a swift adaptive response without compromising essential protein functions.

Building on evidence that subgenomes can diverge functionally in tetraploids (*53*, *70*), we conducted GO enrichment analyses of biased genes across four key tissues (eye, brain, head kidney, and skin) of the cavefish *S. tianlinensis*. This uncovered a consistent pattern of functional partitioning. Genes biased toward the D subgenome were enriched in active and adaptive processes, including tissue remodelling (eye), neural plasticity (brain), and heightened immune responses (kidney and skin). Conversely, genes biased toward the S subgenome were consistently enriched in core housekeeping functions such as basal energy metabolism and maintaining organ and barrier integrity (Supplementary Fig. 10a–h). This suggests that the D subgenome plays a disproportionate role in adaptive responses to cave environments, while the S subgenome mainly maintains essential physiological processes.

The enrichment of adaptive functions on the D subgenome shows that subgenomes can take on different evolutionary roles after polyploidisation. The D subgenome appears to have established itself as a primary locus of evolutionary innovation following polyploidisation, a role that is conserved across both cave and surface lineages. This asymmetry suggests that the ancestral D subgenome may have harboured a greater reservoir of standing genetic variation or a more versatile regulatory framework. This inherent evolutionary potential was then leveraged for niche-specific adaptations after speciation; for instance, it was co-opted for survival in the novel underground environments encountered by *S. tianlinensis*. Meanwhile, the S subgenome preserved essential housekeeping functions, freeing the D subgenome from pleiotropic constraints and allowing its adaptive gene networks to develop more independently. Such functional segregation likely offered a way to resolve genomic conflicts and dosage imbalances inherent to polyploidy, supporting the long-term evolutionary success of the lineage (*5*, *71*). This division of roles maintains core physiological stability while enabling significant phenotypic adaptation to a new niche.

### Epigenetic Regulation Linked to Cave Adaptation

To characterise the DNA methylation landscapes of *S. tianlinensis* and *S. longibarbatus*, we performed whole-genome bisulfite sequencing on eye and liver tissues from both species. This approach yielded high-quality data with deep coverage and high bisulfite conversion efficiencies (Supplementary Table 23). Consistent with the typical vertebrate pattern, we found that DNA methylation occurred almost exclusively in the CpG context, with weighted whole-genome levels ranging from 79.00% to 81.20% across all samples. In contrast, we observed negligible non-CpG methylation (both CHG and CHH) at levels below 1.5% (Supplementary Table 24). Based on these findings, we focused all subsequent analyses solely on CpG methylation.

A prevailing hypothesis proposes that subgenome dominance, observed through biased gene retention and expression, might be driven by epigenetic modifications such as DNA methylation (*6*, *72*, *73*). To investigate this in *Sinocyclocheilus*, we examined CpG methylation (mCG) levels across 16,626 homoeologous protein-coding genes and their 2-kb flanking regions in eye and liver tissues of both species. Our analysis revealed no significant and consistent differences in CpG methylation between homoeologous gene pairs in either species or tissue type (Figs. 4a–d). This finding is consistent with previous studies on several tetraploid cyprinids, which reported no significant differences in CG methylation between homoeologous genes (*54*). Similarly, in goldfish, subgenomes exhibit indistinguishable levels of DNA methylation in both gene bodies and promoter regions across multiple tissues and developmental stages (*55*).

**Fig. 4.**
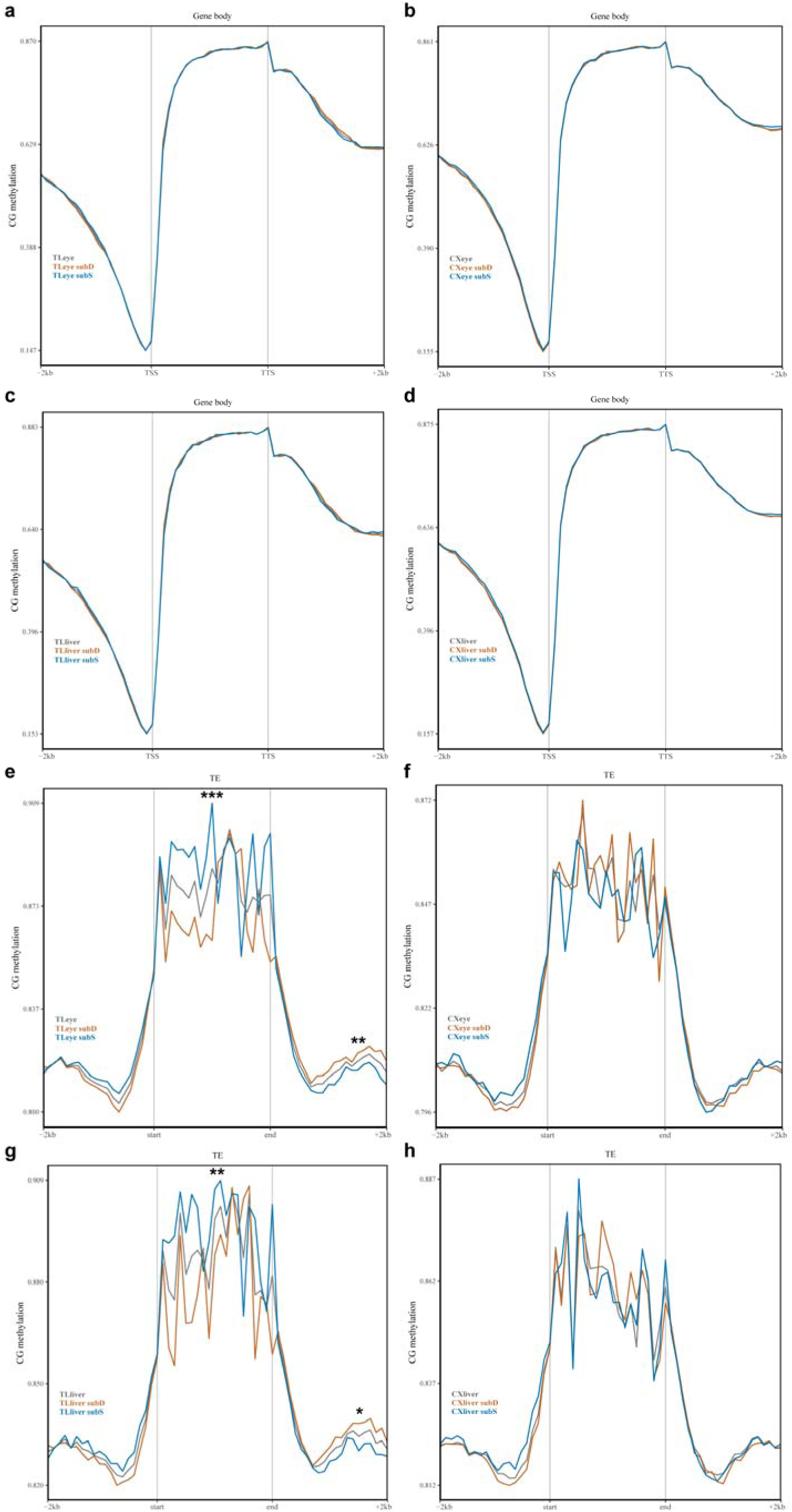
CG methylation levels in subgenomes D and S of eye and liver tissues from *S. tianlinensis* and *S. longibarbatus*. *, P-value <D**0.05 (Wilcoxon rank-sum test); **, P-value <**D**0.01 (Wilcoxon rank-sum test); ***, P-value <**D**0.001 (Wilcoxon rank-sum test).** Panels a-d: Methylation of gene bodies of homoeologous genes and their 2 kb flanking regions. (a) Eye tissue of *S. tianlinensis*. (b) Eye tissue of *S. longibarbatus*. (c) Liver tissue of *S. tianlinensis*. (d) Liver tissue of *S. longibarbatus*. Panels e-h: Methylation of transposable elements (TEs) within 1 kb of homoeologous genes and their 2 kb flanking regions. (e) Eye tissue of *S. tianlinensis*. (f) Eye tissue of *S. longibarbatus*. (g) Liver tissue of *S. tianlinensis*. (h) Liver tissue of *S. longibarbatus*.

A distinct, species-specific pattern was observed when analysing transposable elements (TEs) located within 1 kb of these homoeologous genes. In the cave ecotype *S. tianlinensis*, significant differences were found (Figs. 4e, 4g). TE bodies associated with subgenome S showed higher mCG levels than those associated with subgenome D (Wilcoxon rank-sum test, P = 0.00096 and 0.00580 for eye and liver, respectively), while their downstream regions exhibited significantly lower mCG levels (P = 0.00803 and 0.01491). Conversely, no significant differences in TE methylation between subgenomes were detected in the surface ecotype *S. longibarbatus* (Figs. 4f, 4h; P > 0.05).

These results suggest that asymmetric TE methylation is a lineage-specific characteristic of the cave ecotype *Sinocyclocheilus*. A primary adaptive driver for this change may be the strengthened selection for efficient TE silencing in the energy-limited cave environment. TE transposition is an energetically costly process, and its stringent suppression via DNA methylation would offer a significant energy-saving advantage in this resource-poor habitat. Beyond this direct silencing effect, such differential methylation of TEs could also serve as an evolutionary substrate for regulatory innovation. Modifying the local epigenetic landscape may have contributed to the reorganisation of adjacent gene expression networks, offering additional genomic flexibility to adapt to subterranean life (*56*).

To investigate functional divergence between the subgenomes of the cave ecotype *S. tianlinensis*, we identified homoeologous gene pairs that were differentially methylated specifically within the gene-body. Functional annotation of these pairs showed a distinct pattern of role partitioning that is highly consistent with the known functions of gene-body methylation. Genes with higher gene-body methylation in the D subgenome were enriched in active and specialised processes, including eye development, immune responses, and signal transduction. Conversely, genes with higher gene-body methylation in the S subgenome were mainly associated with essential housekeeping functions, such as basal energy metabolism and the synthesis of core cellular components (Supplementary Fig. 11; Supplementary Table 26). This aligns with the established role of high gene-body methylation in marking stably and broadly expressed genes. These findings indicate that differential gene-body methylation contributes to a division of labour between the two subgenomes. This arrangement may represent a polyploid-specific evolutionary strategy, where one subgenome (S) acts as a stable repository for vital functions, and the other (D) functions as a more adaptable toolkit for evolving specialised traits. Such a division could offer a significant advantage by maintaining the robustness of core biological processes while allowing adaptive flexibility.

This functional partitioning is particularly relevant given the selective pressures faced by *S. tianlinensis* in its cave habitat. The enrichment of eye-development genes among differentially methylated loci in the D subgenome suggests that epigenetic regulation is intimately associated with stygomorphic trait evolution. While it is challenging to disentangle cause and effect—as differential methylation could be a consequence of underlying genetic changes—these epigenetic modifications at least represent a key molecular signature of eye degeneration, consistent with findings in other cavefish such as Astyanax mexicanus (*74*). By localising the regulation of specialised and metabolically costly developmental pathways within a single subgenome, *S. tianlinensis* may more efficiently modulate their expression under aphotic conditions, while safeguarding essential functions in the S subgenome. This epigenetic plasticity may therefore constitute a key mechanism enabling rapid adaptation to novel or extreme environments.

### Divergence in 3D Chromatin Architecture

To investigate the three-dimensional chromatin architecture of the subgenomes, we analysed Hi-C data to examine A/B compartments, topologically associated domains (TADs), and chromatin loops in both species. Genomes are partitioned into transcriptionally active (A) and inactive (B) compartments. In both species, we observed a modest enrichment of the A compartment in the D subgenome relative to the S subgenome. In *S. tianlinensis*, the D and S subgenomes comprised 55.53% and 55.15% A compartments, respectively. A similar pattern was observed in *S. longibarbatus*, with 58.69% and 55.13% A compartments (Fig. 5c; Supplementary Table 27). This higher proportion of euchromatic, transcriptionally permissive A compartments in the D subgenome suggests a stronger contribution to the transcriptome, a trend also reported in other allopolyploid systems such as cotton and cyprinids (*54*, *75*).

**Fig. 5.**
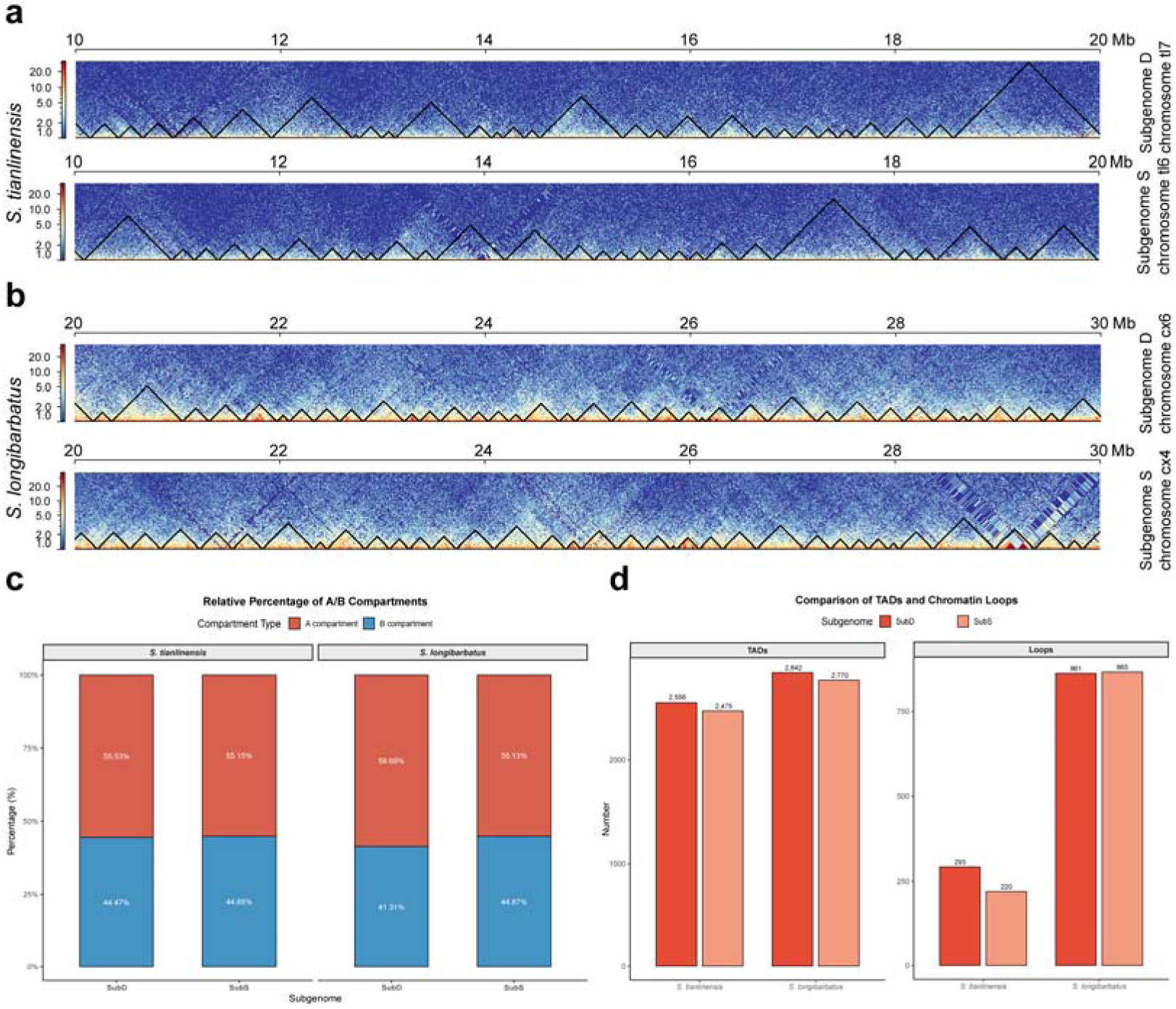
Three-dimensional (3D) genome architectures of subgenomes from *S. tianlinensis* and ***S. longibarbatus*.** (a, b) Hi-C interaction heatmaps showing TAD structures on representative homologous chromosomes from (a) *S. tianlinensis* (chromosomes tl7 and tl6) and (b) *S. longibarbatus* (chromosomes cx6 and cx4). Black triangles delineate TADs. The color scale indicates chromatin interaction frequency, from high (red) to low (blue). (c) Relative percentage of A (active) and B (inactive) compartments in Subgenome D and Subgenome S for both species. (d) Number of identified TADs and chromatin loops in Subgenome D and Subgenome S of *S. tianlinensis* and *S. longibarbatus*.

At higher resolution, we identified TADs, detecting 5,031 in *S. tianlinensis* (2,556 in SubD; 2,475 in SubS) and 5,612 in *S. longibarbatus* (2,842 in SubD; 2,770 in SubS) (Fig. 5d). Examination of TAD boundaries within syntenic regions showed significant shifts between homoeologous chromosome pairs, including tl6/tl7 in *S. tianlinensis* and cx4/cx6 in *S. longibarbatus* (Figs. 5a, 5b). Such boundary shifts can reposition homoeologous genes into new regulatory neighbourhoods, altering their exposure to enhancers or silencing elements and thus providing a structural mechanism for functional divergence (*76*).

Finally, we identified chromatin loops that facilitate long-range transcriptional regulation (*77*). We detected 513 loops in *S. tianlinensis* (293 in SubD; 220 in SubS) and a substantially larger number, 1,726, in *S. longibarbatus* (861 in SubD; 865 in SubS) (Fig. 5d). The near-symmetrical distribution of loops between the subgenomes in *S. longibarbatus* suggests that, while an initial asymmetry between subgenomes was established early, subsequent evolution in this species has involved a global amplification of regulatory connections that transcended the original bias.

### Genomic Loci Associated with Ecotype Differentiation

To identify genetic loci associated with the repeated evolution of cave- and surface-dwelling ecotypes, we performed an association analysis across a broad phylogenetic spectrum of 22 *Sinocyclocheilus* species, including 72 individuals from 10 cave and 12 surface species. We acknowledge that this cross-species design presents strong phylogenetic confounding, and thus our analysis should be interpreted as a scan for candidate loci potentially under parallel or divergent selection, rather than a formal GWAS.

To mitigate the strong effect of shared ancestry, we employed a mixed linear model (LMM), which incorporates a kinship matrix to correct for population structure. This analysis identified 2,406 loci significantly associated with the ecotype (-log10P > 7.71). These loci were distributed across both subgenomes, with 1,250 located in subgenome D and 1,156 in subgenome S (Supplementary Fig. 12). Annotation of these significant loci revealed numerous candidate genes previously linked to traits characteristic of cave adaptation. Notable examples include genes involved in eye development (CRYAB, LMX1B, OPN5, GJA1, OLFM2, PAX6, KERA), pigmentation (TYR, TYRP1), metabolic adaptation (HIF1A, PPARA, MTOR, GHSR), nervous system function (CLOCK), and rhythmic behaviour (BDNF).

These results indicate that adaptation to subterranean environments is a polygenic process, shaped by the coordinated action of multiple loci that influence both regressive traits, such as eye and pigmentation loss, and constructive traits, including metabolic and sensory adaptation.

## CONCLUSIONS

Our study shows that the diversification of *Sinocyclocheilus* is not the result of repeated rounds of polyploidisation, but rather reflects the long-term consequences of a single ancient duplication and the evolutionary processes that followed it. We demonstrate that all species in the genus share a common allotetraploid origin derived from the carp-specific fourth whole-genome duplication (CS4R), with no subsequent ploidy accumulation. This event, which was also shared with related lineages such as *Cyprinus* and *Carassius*, created a state of extensive genomic redundancy and potential. The extraordinary adaptive diversification of *Sinocyclocheilus* unfolded not through additional duplications, but by leveraging the evolutionary potential provided by two key features of its polyploid genome: extensive chromosomal reorganisation and an established pattern of asymmetric evolution between its two subgenomes.

Chromosome-level assemblies of *S. tianlinensis* and *S. longibarbatus* reveal that chromosomal rearrangements, such as fusions, have contributed to karyotype evolution within the genus, giving rise to distinct lineages with diploid chromosome numbers of 2n = 100 and 2n = 96. These events are characteristic of diploidisation and provide a structural mechanism for reproductive isolation and accelerated cladogenesis.

Within this structural framework, we identify a consistent pattern of subgenome dominance. The D subgenome retains more genes, expresses them at higher levels, and occupies a slightly greater fraction of active A compartments in 3D nuclear space. It also harbours more duplications, which are a recognised source of evolutionary novelty. Conversely, the S subgenome has undergone greater gene loss and appears to function primarily as a repository for essential housekeeping functions. This division of labour is reinforced by tissue-specific enrichment of adaptive functions on the D subgenome, particularly in cave populations, where genes related to neural remodelling, immune function, and sensory plasticity are disproportionately represented.

Although global CpG methylation patterns do not differ significantly between subgenomes, we uncover a striking cave-specific signal in the methylation of TEs. In *S. tianlinensis*, but not in *S. longibarbatus*, TEs associated with the S subgenome are hypermethylated relative to their D subgenome homoeologues. This asymmetric regulation likely modulates the local regulatory environment, reshaping expression networks without necessitating widespread changes in coding sequence. Differentially methylated homoeologous pairs further underscore this partitioning, with D-biased genes enriched for costly or specialised processes such as eye development and immune responses, and S-biased genes for metabolic housekeeping. This pattern suggests that the D subgenome has become a dynamic toolkit for adaptation, while the S subgenome maintains physiological stability.

Our research combines genome-wide selection signals with functional genomic data to provide an integrated understanding of cave adaptation. Evolution in subterranean environments demonstrates how natural selection responds to new environmental challenges, eliminating unnecessary traits under relaxed selection while promoting new ones through positive selection. More broadly, our study explores the genetic basis of a major evolutionary transition and provides a framework for understanding how organisms adapt to extreme environments more broadly. These results suggest a model where the CS4R duplication supplied the genomic raw material, large-scale chromosomal rearrangements created structural opportunities for speciation, and the evolution of subgenome dominance, strengthened by epigenetic mechanisms, enabled precise adaptation to subterranean environments. Finally, these genomic, structural, and regulatory processes have driven the remarkable diversification of *Sinocyclocheilus*, establishing it as a key vertebrate model for examining both the evolutionary effects of polyploidy and the mechanisms of ecological specialisation.

## MATERIALS AND METHODS

### Overview of experimental and analytical framework

We combined cytogenetic, genomic, transcriptomic, methylomic, and comparative evolutionary analyses to characterise genome architecture, subgenome evolution, and regulatory asymmetry in *Sinocyclocheilus tianlinensis* (eyeless) and *S. longibarbatus* (normal-eyed). The workflow comprised specimen sampling and karyotyping; Illumina short-read, PacBio HiFi, Hi-C, RNA-seq, and whole-genome bisulfite sequencing; genome assembly and chromosome-scale scaffolding; repeat and gene annotation; comparative analyses of genome evolution, whole-genome duplication (WGD), and synteny; tests for positive and relaxed selection; subgenome identification and validation; detection of structural variation and homoeologous exchanges; expression and methylation asymmetry analyses; inference of 3D genome architecture; and and an association analysis for ecotype-differentiated loci..

### Sampling and karyotype analysis

Two adult females of *S. tianlinensis* (Tianlin County, Guangxi, China) and *S. longibarbatus* (Libo County, Guizhou, China) were sampled under the Laboratory Animal Welfare Ethics Committee of Guangxi University approval (GXU2024279). All *S. tianlinensis* individuals used in this study were maintained under constant aphotic (0L:24D) conditions for at least four weeks prior to sampling to simulate their natural cave environment. For each species, one individual was anaesthetised with MS-222 and dissected. For *S. tianlinensis*, brain, eye, gill, heart, kidney, liver, muscle, skin, and spleen were collected for RNA-seq; for *S. longibarbatus*, brain, eye, gill, heart, swim bladder, liver, muscle, and spleen were sampled. Eye and liver from these individuals were also used for whole-genome bisulfite sequencing (WGBS). All tissues were flash-frozen in liquid nitrogen and stored at −80 °C. A second individual per species provided fresh head kidney tissue for karyotyping.

Karyotyping followed colchicine pretreatment (0.01% for 4 h), hypotonic incubation in 0.075 mol/L KCl for 50 min, and fixation in ice-cold Carnoy’s solution (3:1 methanol:acetic acid). Fixed tissue was macerated in 50% acetic acid, dropped onto pre-warmed slides, air-dried, and stained with 10% Giemsa. At least 50 metaphase spreads per specimen were examined, and representative spreads were imaged using a high-resolution digital system.

### Genome sequencing and Hi-C sequencing

Genomic DNA was extracted from muscle tissue using a QIAamp DNA Mini Kit (Qiagen, USA). A paired-end library with a 350 bp insert size was constructed using the TruSeq Nano DNA HT Sample Preparation Kit (Illumina, USA) and sequenced on an Illumina NovaSeq 6000 platform to generate 150 bp reads. After filtering raw data to remove adapter sequences and low-quality reads, we performed a genome survey analysis on the clean reads. The genome size, heterozygosity rate, and repeat content were estimated using k-mer analysis with the GCE v1.0.2 software (*78*), setting the k-mer size to 17.

For HiFi read generation, DNA fragments >30 kb were selected using the BluePippin System (Sage Science, USA). The library was prepared using the SMRTbell Template Prep Kit 2.0 (PacBio, USA) and sequenced in Circular Consensus Sequencing (CCS) mode on the PacBio Sequel II platform.

Hi-C libraries from muscle followed the protocol of Belton and colleagues with minor modifications (*79*). Chromatin was cross-linked with 4% formaldehyde, nuclei were lysed, and DNA was digested with MboI. Overhangs were filled with biotinylated nucleotides and ligated under proximity-favouring conditions, followed by purification, shearing, and streptavidin capture of biotin-labelled junctions. Libraries were sequenced on an Illumina NovaSeq.

This sequencing effort yielded a comprehensive dataset for two species (Supplementary Table 2). For *S. tianlinensis*, we generated 94.87 Gbp of PacBio HiFi reads (approximately 46.37-fold coverage), 86.05 Gbp of Illumina short reads (approximately 42.06-fold coverage), and 174.79 Gbp of Hi-C data (approximately 85.43-fold coverage). For *S. longibarbatus*, we obtained 66.20 Gbp of HiFi reads (approximately 35.23-fold coverage), 101.72 Gbp of Illumina reads (approximately 54.14-fold coverage), and 201.26 Gbp of Hi-C data (approximately 107.11-fold coverage).

### RNA sequencing

Total RNA was extracted from the previously collected tissues using TRIzol reagent (Invitrogen, USA). Strand-specific mRNA sequencing libraries were then constructed with the Illumina TruSeq Stranded mRNA Library Prep Kit according to the manufacturer’s instructions. The libraries were sequenced on the Illumina NovaSeq 6000 platform (Novogene, Beijing, China). Raw reads underwent a stringent filtering process to remove adapter sequences, reads with poly-N content, and low-quality reads to obtain high-quality data for analysis.

### Whole-genome bisulfite sequencing (WGBS)

WGBS was performed on eye and liver tissues from both *S. tianlinensis* and *S. longibarbatus*. Genomic DNA was extracted using a HiPure Tissue DNA Mini Kit (Magen, China), and its quality and concentration were confirmed via agarose gel electrophoresis and Qubit fluorometry, respectively. For library construction using the Accel-NGS Methyl-Seq DNA Library Kit (Swift, USA), DNA was first fragmented to 200-400 bp with a Covaris S220 instrument. The fragments then underwent bisulfite treatment, which converts unmethylated, but not methylated, cytosines to uracils. After conversion, adapters were ligated, and the library was amplified via PCR. The final library quality was validated on an Agilent 5400 system and quantified by qPCR before being subjected to paired-end sequencing on an Illumina platform.

### Genome assembly

Initial, contig-level assemblies for *S. tianlinensis* and *S. longibarbatus* were generated from the PacBio HiFi reads using Hifiasm v.0.25.0 with default parameters (*80*). Following the initial assembly, we used purge_dups v1.2.6 (*81*) to remove duplicate contigs resulting from high heterozygosity. The assembly was then polished with NextPolish2 using default parameters (*82*). Previous studies on the *Sinocyclocheilus* genome have shown that assemblies generated with this pipeline closely match flow cytometry-based genome size estimates, with a discrepancy of only ∼2% (*32*). To upgrade the assemblies to the chromosome level, we performed Hi-C-based scaffolding. First, quality-controlled Hi-C reads were mapped to the purged primary assembly using BWA-MEM v0.7.19 with the -5SP option (*83*). The resulting alignments were then processed to remove PCR duplicates with samblaster and filtered using samtools view -F 3340 to retain only high-quality, primary alignments (*84*). These preprocessing steps followed the recommended pipeline for HapHiC v1.0.6, a tool we selected based on evaluations showing its superior performance over other Hi-C scaffolders (*85*). The filtered alignments were then used to scaffold the contigs into chromosomes using HapHiC with default parameters. The target chromosome number was set to 50 for both species, consistent with their known tetraploid karyotypes (2n=100). The interaction matrices were converted to .hic format using the juicer_post utility provided with HapHiC, and manual inspection and correction of contig order and orientation were performed in Juicebox v2.17.0 (*86*). The final assembly was assessed using BUSCO (Benchmarking Universal Single-Copy Orthologs) with the Actinopterygii_odb10 lineage database (*87*).

### Repeat annotation

The identification and annotation of repetitive elements in the *S. tianlinensis* and *S. longibarbatus* genomes were performed using the earlGrey v6.1.1 pipeline (*88*). This pipeline integrates several tools to create a comprehensive and curated repeat library. The earlGrey workflow proceeded as follows. First, known repetitive elements were identified by masking the genome with RepeatMasker v4.1.8 against the Dfam database v3.8 (*89*). Second, de novo repeat discovery was performed using RepeatModeler v2.0.4 (*90*). The elements identified from both approaches were then combined, processed to remove redundancy, and used to generate an optimized set of consensus sequences through an automated "BLAST, extract, extend" protocol. Finally, the genome was re-masked with RepeatMasker using this newly curated, custom library. The results were combined with the output from LTRfinder and further post-processed with RepeatCraft to generate the final repeat annotation and landscape (*91*, *92*).

### Gene structure and function annotations

Protein-coding gene prediction was performed using a combined approach of homology-based, ab initio, and RNA-Seq-assisted predictions with the BRAKER3 pipeline v3.0.8 (*93*). First, transcriptomes derived from nine tissues of *S. tianlinensis* (brain, eye, gill, heart, kidney, liver, muscle, skin, and spleen) and eight tissues of *S. longibarbatus* (brain, eye, gill, heart, swim bladder, liver, muscle, and spleen), generated in this study, were mapped to the soft-masked genome using HISAT2 v2.1.1 (*94*). These transcriptome assemblies (BAM files) provided RNA evidence for BRAKER3, which utilized both AUGUSTUS v3.5.0 and GeneMark-ETP v1.0.2 to refine gene predictions by training models on RNA-Seq-derived evidence, improving gene structure accuracy (*95*, *96*). For homology-based evidence, a comprehensive protein database was constructed by combining protein sequences from the OrthoDB v11 Vertebrata database with those from eight related species (*S. jii*, *S. rhinocerous*, *S. anshuiensis*, *S. grahami*, *S. maitianheensis*, *S. anophthalmus*, *Cyprinus carpio*, and *Carassius auratus*). This database was used to enhance annotation through protein sequence alignment and homology-informed model training in AUGUSTUS. Ab initio prediction was performed using models trained on RNA-Seq and protein evidence data, enabling BRAKER3 to infer gene structures with minimal manual intervention. The results from AUGUSTUS and GeneMark-ETP were combined using TSEBRA v1.1.2 to produce the final gene predictions (*97*). Finally, the PASA v2.5.3 pipeline was used to update the structural annotation, incorporating untranslated regions (UTRs) and improving gene-isoform relationships (*98*). The completeness of the annotated gene sets for both species was evaluated with BUSCO against the actinopterygii dataset.

To functionally annotate the predicted genes, we performed homology searches of the protein sequences against the NCBI non-redundant (Nr), SwissProt, and TrEMBL databases using BLASTP (E-value < 1e-5) (*99*). Protein domains and families were identified by searching against the InterPro database. Furthermore, Gene Ontology (GO) terms and Kyoto Encyclopedia of Genes and Genomes (KEGG) pathways were assigned using eggNOG-mapper based on orthology mapping (*100*).

### Genome evolution

To investigate the genomic evolution of *S. tianlinensis* and *S. longibarbatus*, we performed a comparative analysis including all available *Sinocyclocheilus* genomes (*S. jii*, *S. rhinocerous*, *S. anshuiensis*, *S. grahami*, *S. maitianheensis*, *S. anophthalmus*) and two outgroup species (*Cyprinus carpio* and *Carassius auratus*). All of these species are of tetraploid origin.

We used OrthoFinder v3.0.1 with default parameters to identify orthogroups (*101*). A species tree was inferred from 3,710 single-copy orthologs using the STAG algorithm and rooted with STRIDE, with STAG support values at internal nodes (*102*). This filtering strategy isolates a consistent set of genes that have likely returned to a single-copy state following the shared whole-genome duplication event, thus providing a robust dataset for phylogenetic inference. To analyze gene family dynamics, we used CAFE v5.1.0 (*103*). The species tree was first made ultrametric using the make_ultrametric.py script from OrthoFinder. We then ran CAFE with a Poisson distribution to generate an error model, which was subsequently used in three replicate runs to ensure convergence. Gene families with a family-wide P-value < 0.05 were considered to have undergone significant expansion or contraction. Finally, Gene Ontology (GO) enrichment analysis was performed on genes from these significantly changed families in the four *Sinocyclocheilus* cave ecotype species (*104*).

### Whole-genome duplication

We investigated whole-genome duplication (WGD) events in *S. tianlinensis* and *S. longibarbatus* using the WGDI v0.7.1 pipeline (*105*). For comparative context, the analysis included six other *Sinocyclocheilus* species (*S. jii*, *S. rhinocerous*, *S. anshuiensis*, *S. grahami*, *S. maitianheensis*, *S. anophthalmus*) and two outgroup species (*Cyprinus carpio* and *Carassius auratus*).

First, an all-versus-all BLASTP search (E-value < 1e-5) was performed within each genome to identify intraspecific homologous gene pairs. These homologs were then processed with WGDI (-icl parameter) to detect collinear blocks. Synonymous (Ks) and non-synonymous (Ka) substitution rates for these collinear gene pairs were calculated within WGDI (-ks parameter), which implements the codeml program from the PAML v4.9 package (*106*). Collinear blocks were subsequently partitioned based on their Ks distributions (-kp parameter), and the resulting distribution was used to fit Ks peaks (-pf parameter), with each peak representing the median Ks of a set of homologous blocks. The age (T) of each WGD event was estimated using the formula T = Ks / (2r), where Ks is the median value of the fitted peak and r is the neutral substitution rate, assumed here to be 3.51 × 10[[substitutions per synonymous site per year (*107*).

### Genome synteny

We analyzed syntenic relationships among the ten fish genomes using the MCscan toolkit from JCVI (*108*). First, GFF3 annotation files for all species (*S. jii, S. tianlinensis, S. rhinocerous, S. anshuiensis, S. longibarbatus, S. grahami, S. maitianheensis, S. anophthalmus, Cyprinus carpio,* and *Carassius auratus*) were converted to BED format. Syntenic blocks were then identified through a series of pairwise comparisons designed to trace chromosomal evolution sequentially through the phylogeny (e.g., *Carassius auratus* vs. *Cyprinus carpio*, *Cyprinus carpio* vs. *S. jii*, etc.), using the python3 -m jcvi.compara.catalog ortholog command. The resulting pairwise synteny and large-scale macrosynteny were visualized using the jcvi.compara.synteny and jcvi.graphics.karyotype modules, respectively.

### Analyses of positive and relaxed selection

To investigate the molecular evolution of protein-coding genes, particularly those potentially driving adaptation in cavefish, we scanned for signatures of positive selection. The analysis was based on the 3,710 single-copy orthologs identified previously. First, we established a robust alignment for each orthogroup. We used PRANK v.170427 to generate codon-based alignments, leveraging the species tree as a guide to improve accuracy (*109*). Ambiguous or poorly aligned sites were subsequently removed from these alignments using Gblocks v0.91b (*110*). With these filtered alignments, we then tested for positive selection using the branch-site model within the codeml program of PAML4, executed via the GWideCodeML v1.1 wrapper (*111*). This model is specifically designed to detect episodic positive selection on particular branches of a phylogeny. We designated the four cave ecotype *Sinocyclocheilus* lineages as the ’foreground’ to test for selection events associated with their adaptation to subterranean life. A gene was considered a candidate for positive selection only if it met a stringent set of conditions, which were filtered using a custom Python script. These conditions included: an estimated ω (the ratio of nonsynonymous to synonymous substitution rates, dN/dS) greater than 1 on the foreground lineages, a significant likelihood ratio test (P < 0.05) supporting the positive selection model over a null model, and a high-quality alignment spanning at least 150 amino acid residues.

To determine if adaptation to the cave environment involved a relaxation of selective constraints on certain genes, we employed the RELAX model (*112*), implemented in the HyPhy v.2.5.8 package (*113*). This analysis was performed on the full set of 3,710 single-copy orthologs. The RELAX method explicitly tests for changes in selection strength by comparing a designated ’test’ (foreground) set of branches to the remaining ’reference’ (background) branches in a phylogeny. We defined the four cavefish lineages as the foreground for this test. The model works by fitting a distribution of selection intensities (ω) to the reference branches and then assessing whether this distribution is shifted for the foreground branches via a selection intensity parameter

(K). A K value < 1 indicates that the selective strength has been relaxed (e.g., purifying selection has weakened or positive selection has diminished), while K > 1 suggests an intensification of selection. The null model, which posits no change in selection intensity, is represented by K=1. We tested for significant deviations from this null model using a likelihood ratio test (LRT). Genes were classified as evolving under relaxed selection if the alternative model provided a significantly better fit (P < 0.05 after Holm-Bonferroni correction) and the estimated K value was less than 1. The sets of genes identified as being under positive and relaxed selection were then subjected to Gene Ontology (GO) enrichment analysis.

### Subgenome identification

We partitioned the *S. tianlinensis* and *S. longibarbatus* genomes into their constituent subgenomes (subD and subS) using the SubPhaser pipeline (*52*). First, we generated 15-mer frequency counts for each chromosome using Jellyfish v2.2.6 (*114*). Next, divergent k-mers were identified between homoeologous chromosome pairs and clustered into two subgenomes using a K-means algorithm. The final phasing was refined and validated through hierarchical clustering and principal component analysis (PCA). This approach has been successfully applied to phase the subgenomes of other polyploid cyprinids, including *Cyprinus carpio*, *Procypris rabaudi*, *Spinibarbus sinensis*, and *Luciobarbus capito* (*52*, *54*).

### Phylogenetic tree reconstruction of subgenomes

Our comparative genomic analysis included the genomes of 17 cyprinid species and the zebrafish (*Danio rerio*) as an outgroup. The dataset comprised seven diploid species (*Cirrhinus molitorella*, *Ctenopharyngodon idella*, *Onychostoma macrolepis*, *Paracanthobrama guichenoti*, *Poropuntius huangchuchieni*, *Puntius tetrazona*) and ten tetraploid species. For the tetraploid species (*Cyprinus carpio*, *Carassius auratus*, and the eight *Sinocyclocheilus* species), we analyzed their constituent D and S subgenomes separately, which were assigned based on synteny with our reference *S. tianlinensis* and *S. longibarbatus* subgenomes. Using this set of 27 genomes (20 subgenomes + 7 diploid genomes) plus the outgroup, we identified single-copy orthogroups using OrthoFinder with default parameters.

From the OrthoFinder results, a set of single-copy orthologs present in all species was selected. For each orthogroup, the protein sequences were aligned using MAFFT v7.453 (*115*), and the protein alignments were converted to codon alignments using pal2nal v14.1 (*116*). These individual codon alignments were then concatenated into a supermatrix. This supermatrix was used to construct a maximum likelihood (ML) species tree with IQ-TREE v2.2.6 (*117*). The analysis was run with 1,000 bootstrap replicates, and the best-fit substitution model was determined by ModelFinder (*118*). Subsequently, divergence times were estimated on the ML topology using the penalized likelihood method implemented in treePL (*119*). The dating analysis was calibrated with the following three-time constraints: (1) the divergence of *Danio rerio* from the common ancestor of common carp and goldfish: 32.81–36.84 Mya (*56*), (2) the divergence between the D subgenomes of *Cyprinus carpio* and *Carassius auratus*: 8.14–9.14 Mya (*53*), (3) the divergence between the S subgenomes of *Cyprinus carpio* and *Carassius auratus*: 7.74–8.69 Mya (*53*).

### Chromosomal structure variations and homeologous exchanges (HE) of subgenomes

We identified structural variations (SVs) such as inversions, translocations, and duplications between the homoeologous chromosomes of the subgenomes using SYRI v1.7.1 (*120*). The workflow began with a whole-genome self-alignment using MUMmer v4.0 (*121*). Specifically, we used nucmer with optimized parameters (-maxmatch -c 100 -b 500 -l 50) to generate alignment blocks. These blocks were subsequently filtered for high-confidence, one-to-one alignments using delta-filter (-m -i 90 -l 100). The filtered alignment file was then provided as input to SYRI, which was run with its default parameters to annotate the full spectrum of SVs.

We identified homoeologous exchange (HE) regions between the subD and subS subgenomes following a previously established methodology integrating synteny and read-depth analysis (*122*, *123*). First, syntenic blocks between the subgenomes were established by aligning them with Minimap2 v2.1 and chaining regions separated by less than 20 kb (*124*). Concurrently, Illumina paired-end reads from *S. tianlinensis* and *S. longibarbatus* were separately mapped to their respective chromosome-level assemblies using BWA-MEM2 v2.2.1 (*125*). After filtering secondary alignments with SAMtools, we calculated read depth in 1 kb non-overlapping windows using Bamdst. Windows exhibiting outlier depths—defined as 1.5–5.0x (potential duplications) or 0–0.5x (potential deletions) of the chromosome’s average depth—were identified. Finally, adjacent windows of the same type were merged. A merged region was designated as a putative HE event only if it exceeded 60 kb in length and was located within a pre-identified syntenic block.

### Asymmetric evolution of *S. tianlinensis* and *S. longibarbatus* subgenomes

Recent studies have reported Ohnolog retention bias in certain gene sets, including BUSCO genes, towards one subgenome in species such as the Prussian carp, goldfish, common carp, *Luciobarbus capito*, *Procypris rabaudi*, and *Spinibarbus sinensis* (*54*, *126*). Building on this, we utilized BUSCO with the Actinopterygii_odb10 lineage database to evaluate this phenomenon for the D and S subgenomes of *Sinocyclocheilus tianlinensis* and *Sinocyclocheilus longibarbatus*.

To further investigate the asymmetric evolution of the D and S subgenomes, we analyzed the differential expression of homoeologs in *S. tianlinensis* and *S. longibarbatus*. For this purpose, clean RNA-seq reads from nine tissues of *S. tianlinensis* (brain, eye, gill, heart, kidney, liver, muscle, skin, and spleen) and eight tissues of *S. longibarbatus* (brain, eye, gill, heart, swim bladder, liver, muscle, and spleen) were separately mapped onto their respective genomes using Hisat2 with default parameters. Gene expression levels were quantified as transcripts per million (TPM) using StringTie v2.1.5 (*127*). Homoeolog expression bias (HEB) analysis was conducted on syntenic gene pairs between the subgenomes. These syntenic pairs were identified using the Python version of MCScan within JCVI (parameters: --cscore=.99). After filtering out gene pairs with TPM values below 1 in all samples, pairs with a greater than two-fold difference in TPM were classified as expression-biased, exhibiting either SubD or SubS dominance. Within each biased pair, the gene with the higher expression level was designated as the dominant gene (DG), while the other was termed the submissive gene (SG). Genes in pairs without significant expression differences were considered neutral.

### MethylC-seq data analysis

Raw sequencing reads from eye and liver tissues of *S. tianlinensis* and *S. longibarbatus* were first processed with Trim Galore v0.6.10 to trim low-quality bases and remove adapters (*128*). The trimmed reads were then processed using the Bismark v0.24.2 pipeline (*129*). This involved mapping reads to the respective reference genomes, removing PCR duplicates to mitigate amplification bias, and extracting methylation calls (in CG, CHG, and CHH contexts) with the bismark_methylation_extractor. The methylation level for each cytosine was calculated as the ratio of methylated reads to the total read depth at that site.

Metaplots of DNA methylation levels were generated for homologous protein-coding genes and their neighboring transposable elements (TEs) annotated in the *S. tianlinesis* and *S. longibarbatus* genomes. For each gene and TE, weighted DNA methylation levels were calculated across three distinct regions: the 2-kb region upstream of the gene or TE, the gene body (defined as the coding sequence) or TE body, and the 2-kb region downstream. Each of these regions was subsequently divided into 20 bins. The weighted DNA methylation level was then calculated for each bin, and the results were combined to generate metaplots of methylation across the gene/TE body and its flanking regions. To compare methylation levels between the D and S subgenomes, a Wilcoxon rank-sum test was performed for each of the three regions (upstream, body, and downstream).

To determine the methylation status of individual cytosine loci, a binomial test was employed. Specifically, the binom_test function from the SciPy package in Python was utilized to calculate the p-value for each cytosine locus (*130*). This test assesses the probability of observing at least k methylated reads in n total reads, given a predefined error rate p. The parameters were defined as follows: k represents the number of reads supporting a methylated state, n is the total read coverage at that locus, and p is the background probability of methylation, modeled as the bisulfite non-conversion rate or sequencing error. Gene-level methylation was subsequently quantified using an in-house script, which calculated the weighted average methylation for each gene body. To investigate differential methylation between the two subgenomes, the methylation levels of homoeologous gene pairs were compared. A gene pair was classified as differentially methylated if it met two criteria: (1) at least one gene was identified as either hypermethylated (methylation level > 90%) or hypomethylated (methylation level < 1%), and (2) the ratio of their methylation levels exceeded two-fold.

### 3D genome architecture identification

To characterize the 3D genome architecture, filtered and trimmed Hi-C reads from *S. tianlinensis* and *S. longibarbatus* were mapped to their respective genomes using BWA-mem (-A1 -B4 -E50 -L0). The subsequent analysis was performed using the HiCExplorer suite v3.7.6 (*131*). First, contact matrices were generated at 10 kb resolution with hicBuildMatrix. Subsequently, A/B compartments were assigned at 100 kb resolution using hicPCA, with compartment identity determined by eigenvalue direction adjusted for gene density. At a finer scale, topologically associating domains (TADs) and chromatin loops were identified at 10 kb resolution using hicFindTADs and hicDetectLoops, respectively. Finally, the resulting TAD structures were visualized with hicPlotTADs.

### Identification of Genetic Loci Associated with Cave-Surface Ecotype Differentiation

To identify genetic loci highly differentiated between cave- and surface-dwelling ecotypes, we performed an association analysis on previously reported RAD-seq data from 72 *Sinocyclocheilus* individuals. Crucially, this dataset encompasses a complex phylogenetic structure, comprising 10 distinct cave-dwelling species and 12 distinct surface-dwelling species (*17*). Given this cross-species design, our approach is best described as a phylogenetically-informed scan for divergent loci, rather than a traditional within-population GWAS. Filtered reads were aligned to the *S. tianlinensis* reference genome using BWA. SNPs were then called using the bcftools v1.12 mpileup and call pipeline, with a minimum mapping and base quality of 20 (*132*). To ensure a high-quality dataset for association testing, we applied a multi-step filtering cascade with bcftools. Initially, variants were retained if they had a quality score (QUAL) > 30, read depth (INFO/DP) at least three times the number of samples, and a missing data fraction < 0.5. Subsequently, we enforced a minor allele frequency (MAF) > 0.05 and a missing data fraction < 0.1. The final VCF file was processed with PLINK v1.9 for quality control and formatting (*133*).

To perform the association analysis while accounting for the profound population and species structure in our dataset, we employed a mixed linear model (LMM) implemented in GEMMA v0.98.3 (*134*). The LMM incorporates a kinship matrix, calculated from the genomic data, as a random effect to mitigate confounding signals arising from shared ancestry and relatedness, thereby minimising false positives. The entire analysis was executed via the vcf2gwas Python API (*135*). The pipeline was supplied with the filtered VCF file, the binary phenotype data (0=surface ecotype, 1=cave ecotype), and the GFF3 annotation file. The results, including Manhattan plots and lists of candidate SNPs highly associated with ecotype, were generated directly by the CMplot package and the API.

## Acknowledgments

We thank the EED lab members for fieldwork and Cheng-Hai Fu for his field assistance. We also thank Jia-Jun Zhou for providing samples and for his field assistance. We gratefully acknowledge the high-performance computing platform at Guangxi University for supporting part of the bioinformatics analyses.

## Author contributions

T.M. and M.M. conceptualised the research; T.M., M.M. and H.S. designed the methodology; Y.L., T.M., M.M., and J.Y. conducted fieldwork; T.M. carried out formal analysis; T.M., M.M., H.S., M.V., L.Y. and S.H. wrote the original draft; T.M. made figures; M.M., L.Y. and S.H. supervised; all authors reviewed and edited the draft. All authors read and approved the final manuscript.

## Data and materials availability

All the data will be made available upon formal acceptance of the paper.

